# Prioritization of Potential Drugs Through Pathway-Based Drug Repurposing and Network Proximity Analysis

**DOI:** 10.1101/2025.06.05.658004

**Authors:** Belén Otero-Carrasco, Lucía Prieto-Santamaría, Alejandro Rodríguez-González

## Abstract

Drug repurposing is an effective strategy to identify novel therapeutic options by leveraging existing drugs with known mechanisms of action and safety profiles. This work introduces a pathway-based drug repurposing approach that combines protein-protein interaction (PPI) networks and transcriptomic data to prioritize candidate drugs. The underlying assumption is that if disease 1 and disease 2 are both associated with the same biological pathway, then drugs known to be effective for disease 2 may be potential treatments for disease 1. To explore these relationships, disease–pathway–drug triplets are constructed. Genes shared between disease 1 and the associated pathway are identified, and their proximity is calculated within the PPI network using a Z-score based on random permutations. Candidate drugs for disease 2 are then analyzed using the Connectivity Map data, assessing whether their up- or down-regulated gene signatures are enriched for the genes contributing to the proximity between disease 1 and the pathway. This integration enables the ranking of candidate drugs based on their potential biological impact on disease 1. Finally, top-ranked drug–disease associations are evaluated through manual curation of the scientific literature to assess existing evidence supporting the proposed repurposing hypothesis.

## I. Introduction

Drug repositioning has emerged as a powerful strategy to accelerate therapeutic discovery by identifying new uses for existing drugs, reducing development costs, and shortening the time to clinical application [1], [2]. Traditional drug discovery follows a “one drug–one target–one disease” paradigm, where compounds are developed to selectively modulate a single molecular target implicated in a specific disease [3]. However, this approach has limitations, as many diseases arise from complex dysregulations across multiple biological pathways rather than isolated molecular defects [4]. Additionally, drug promiscuity—where a single drug interacts with multiple targets—suggests that the one-target model oversimplifies drug mechanisms [5].

To overcome these limitations, pathway-based drug repositioning has gained traction as a more holistic alternative. This approach recognizes that diseases sharing common dysregulated pathways may be treated with similar drugs, even if their primary indications differ [6]. By focusing on pathway-level perturbations rather than individual targets, this strategy captures the polypharmacological nature of drugs and the systemic complexity of diseases [7]. Several computational methods have been developed to exploit pathway-based associations, including gene set enrichment analysis [8], pathway activity scoring [9], and network propagation techniques [10].

A key advantage of pathway-based repositioning is its ability to uncover indirect drug-disease relationships that would be missed by traditional target-centric approaches. For instance, a drug that modulates a pathway upstream or downstream of a disease-associated gene may still exert therapeutic effects without directly binding to the primary disease gene [11]. Protein-Protein Interaction (PPI) networks provide an ideal framework for such analyses, as they enable the systematic evaluation of drug-disease relationships by measuring topological proximity between drug targets and disease modules [12]. Building upon this network-based perspective, Aguirre-Plans *et al*. [13] introduced the concept of Proximal Pathway Enrichment Analysis (PxEA) to identify drugs capable of targeting shared biological processes in comorbid diseases. By quantifying the proximity between drug targets and disease-associated pathways within the PPI network, this method facilitates the identification of repurposing candidates that may modulate key nodes implicated in multiple conditions. Their approach underscored the potential of network endopharmacology to simultaneously address the biological complexity of diseases and the multi-target nature of drugs.

Inspired by these concepts, our study proposes a pathway-centered drug repurposing framework that integrates network proximity analysis with transcriptomic signatures. Specifically, we leverage the Connectivity Map (CMap) resource [14], [15], which provides large-scale gene expression profiles of drug-induced perturbations in human cell lines. By aligning CMap transcriptional responses with PPI-based proximity metrics, we aim to prioritize candidate drugs that are not only topologically close to disease– pathway modules, but also capable of functionally modulating their associated gene expression profiles. This integrative strategy enhances the biological interpretability of drug–disease associations and improves confidence in repurposing predictions.

## II. Materials and methods

### A. Overview

In this study, we present a network-based pathway-based drug repurposing approach that integrates PPI networks and transcriptomic data to prioritize candidate drugs. Our methodology is based on the hypothesis that if *Disease 1* and *Disease 2* are both associated with the same biological pathway, then drugs known to treat *Disease 2* may also be viable candidates for *Disease 1*. To test the hypothesis, the following steps are performed:

- *Disease 1* – *Pathway* – *Disease 2* – *Drug* triplets: identifying shared genes between *Disease 1* and the associated pathway.
- Quantifying gene products proximity within the PPI network using a Z-score derived from degree-preserving permutations, ensuring statistical robustness.
- Analyzing drug-induced gene signatures from the Connectivity Map (CMap) to assess whether candidate drugs modulate the genes contributing to the pathway-disease proximity.
- Ranking drug candidates based on their proximity and biological impact at the gene expression level, followed by a literature review to validate reuse hypotheses.

By combining these methods, we provide a robust and scalable framework that goes beyond the ‘one drug, one target’ paradigm, providing a list of drug candidates for the diseases studied based on biological pathways. The complete source code and all results produced during this study can be accessed at the following GitLab repository^1^.

### B. Data selection and integration

The identification of drug repurposing cases in which biological pathways played a central role—referred to as DREBIOP (Drug REpurposing based on BIOlogical Pathways)—was initially conducted in a previous study [16]. Building upon the insights obtained from that investigation, we developed a novel drug repurposing approach that leverages direct relationships between diseases through shared biological pathways [17]. This approach is based on the hypothesis that if *Disease 1* and *Disease 2* are connected via a common biological pathway, then drugs used to treat *Disease 2* may serve as potential therapeutic candidates for *Disease 1*, due to the shared underlying biological mechanism.

The application of this strategy led to the identification of a new set of drug repurposing candidates requiring further validation. In total, 21,968 disease–drug associations were identified, of which 16,848 represented novel associations (that is, cases in which a drug not previously linked to a given disease was connected to it through a specific biological pathway).

Following this, we focused on the subset of triplets of the form *Disease 1* – *Disease 2* – *Drug* in which the drug was novel for Disease 1. The next step involved filtering these triplets to retain only those cases where more than one candidate drug was associated with the same *Disease 1*. This filtering was essential, as the overarching goal was to enable prioritization by ranking candidate drugs based on their potential relevance for a given disease. From an initial set of 821 triplets involving novel drug–disease associations, 556 triplets were retained that involved multiple drug candidates for the same *Disease 1*.

Finally, a further refinement was applied by selecting only those cases in which *Disease 1* and the associated biological pathway(s) shared at least one gene. This additional criterion was introduced to enhance the biological plausibility of the proposed drug–disease associations. After applying this filter, the dataset was reduced to a final set of 155 triplets.

### C. Network distance

We started with these 155 selected triplets, which had the form *Disease 1* – *Pathway* – *Disease 2*. The next step in the methodology focused on the relationship between *Disease 1*, the associated pathway, and the candidate drug. At this stage, the primary objective was to establish a biologically meaningful connection between *Disease 1* and the *Pathway*, which would enable the prioritization of candidate drugs for repurposing.

To achieve this, a network-based approach was employed to compute the proximity between candidate drug targets and genes in the pathway associated with *Disease 1*. Specifically, the distance between the targets of each candidate drug and the genes involved in the relevant biological pathways was calculated within the context of a biological interaction network. This analysis aimed to identify how functionally close each drug was to the disease-specific biological processes, excluding direct associations based solely on the drug’s known target gene. Following this analysis, a total of 48 unique *Disease 1* – *Pathway* relationships were retained. Cases in which the drug’s target gene was the only gene shared between Disease 1 and the pathway were excluded, as the goal was to explore repurposing opportunities beyond direct target overlap. This filtering ensured that the prioritization of candidate drugs was based on broader biological context rather than on existing target-disease associations.

On the other hand, we used the protein-protein interaction (PPI) network, also known as the interactome, to map interactions between proteins in the cell. These PPIs were compiled from neXtProt [18]. For consistency, we standardized proteins as genes, following the NCBI nomenclature and identifiers’ system. A total of 13,979 genes were considered, with 370,267 interactions between them. Although, conceptually, we are studying interactions between gene products rather than genes themselves, this standardization ensures a uniform coding terminology.

For the obtention of a network-based distance, we relied on a well-known metric commonly used in bioinformatic pipelines: the disease-drug proximity [12]. Originally intended for the measurement of the relationship between disease modules and drug targets in the interactome, this metric takes into account two sets of nodes, computes the Shortest Path Lengths (SPLs) between them, identifies closer connections, and averages them across targets. Afterwards, it derives a Z-score from the previous distance by generating reference distributions corresponding to the expected distances between two randomly selected groups of proteins matching the size and the degrees of the original sets of nodes in the network. This Z-score is the metric named as network-based proximity and its formal definition starts by computing the *closest distance* (*d*_*c*_), between disease module proteins and drug targets:

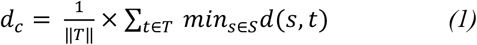

where “*T*” denotes the total protein targets of a drug; “*t*”, each target individually; “*S*”, the set of proteins in the disease module; “*s*”, each protein individually, and “*d*(*s, t*)”, the SPL between each “*s*” and “*t*” pair.

Instead of disease modules, we updated these metrics towards our goal of identifying promising drugs closer to the proteins in the pathways that were associated with *Disease 1* as well. That is, in our case, “*S*” stands for the proteins shared between the disease and the pathway and not for the disease module. For clarity, from this computation process, we report 3 different metrics:

- *Minimum SPL, min*_*t∈T,s∈S*_*d*(*s, t*), which inform us of which are the closer genes in both sets, as well as of the number of hops-away that the drug closest target(s) is from the closest protein(s) in the *Pathway* that are associated to *Disease 1*.
- *Closest distance* or *average SPL, d*_*c*_, which provides the mean of the shortest path distances form nodes in the set of drug targets *T* to the closest nodes in the set of disease-pathway proteins *S*, computed as stated in Equation (1).
- *Proximity*, which measures the Z-score of the previous *closest distance* between the two sets of nodes S and T within the interactome when compared to random expectation. The formula is as follows:

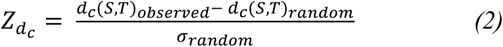

where *d*_*c*_ (*S, T*)_*observed*_ is the actual *closest distance, d*_*c*_ (*S, T*)_*random*_ is the mean of the reference distribution of distances observed if the two sets of nodes are randomly chosen from the interactome (preserving both sets’ degree distributions), and *σ*_*random*_ is the standard deviation of this sampled distance distribution. The obtained 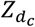 quantifies whether a particular *d*_*c*_ (*S, T*) is smaller than expected by chance.

In summary, for each *Disease* – *Pathway* – *Drug* triplet, we identified proteins codified by shared genes between the *Disease* and *Pathway*, we compiled the target genes of the *Drug*, and we computed the previous 3 metrics: the *minimum SPL*, the *closest distance*, and the *proximity*. During this process, we also identified the specific genes in the two sets that were closer in the network, as well as the genes in the shortest paths to reach one set to the other.

### D. Enrichment analysis

Once the SPLs and proximity metrics between candidate drugs and *Disease 1*-associated pathway gene products were calculated, the next step involved the use of the CMap database to assess whether the genes closest to the disease exhibited enrichment across various cell lines. The goal of this step was to determine whether the gene most proximal to the disease, based on network proximity calculations, was significantly affected by the candidate drug in terms of gene expression modulation.

To this end, we examined whether the expression of the drug-proximal gene was significantly altered upon treatment with the corresponding drug across different cell lines, as reported in CMap. Specifically, the enrichment analysis focused on the Level 5 moderated Z-scores (modZ) provided by CMap. Following established criteria in the scientific literature [15], a gene was considered significantly modulated if the absolute value of its modZ score exceeded 2, indicating a relevant change in expression.

Based on these results, the final step of the methodology consisted of ranking the candidate drugs for each *Disease 1*. This ranking was determined by integrating both the gene enrichment information derived from CMap and the proximity score previously calculated between the drug and the disease-associated pathways. In this way, prioritization accounts for both the network-based biological relevance of the drug and its capacity to modulate disease-related genes at the transcriptional level. Finally, when the most potential drug candidate for each disease studied was obtained, the feasibility of this relationship was validated in scientific literature.

## III. RESULTS AND DISCUSSION

In the present section, we present a detailed description of the results obtained in this study. This section is structured into several subsections, each corresponding to a specific step of the methodology described above. The organization of the results follows the logical progression of our computational pipeline, from the identification of disease–pathway–drug triplets to the prioritization and validation of candidate drug repurposing hypotheses. Given the nature of our pathway-based drug repurposing approach, which integrates PPI networks, transcriptomic signatures, and disease association data, the results are interpreted within the context of biological relevance. The modular structure of the methodology allows for a comprehensive evaluation of each component— proximity analysis within the interactome, enrichment of drug-induced gene expression profiles, and literature-based validation—thereby reinforcing the robustness of the identified candidate drugs.

### A. Data Descriptive Analysis

First, a descriptive analysis was performed on the triplets that form the final dataset studied. This dataset consists of 155 distinct triplets, considering the relationship between *Disease 1, Pathway*, and *Disease 2*. These triplets are composed of a combination of 54 *Disease 1*, 34 *Disease 2*, and 33 *Pathways*. Additionally, a representation of the number of drugs associated with each of these triplets was created (**Figure 1**). The average number of drugs associated with each triplet is 36.57, with a standard deviation of 24.51. Given these results, the prioritization of candidate drugs becomes essential, as proposing such a large number of drugs for a given disease is not the most effective approach to identifying a potential drug for repurposing. The next section will describe the proximity calculation using network medicine to generate an initial ranking of these drugs, thus having a way of prioritizing between the great number of drugs that could be repurposed for each disease as shown in **Figure 1**.

**Fig. 1.**
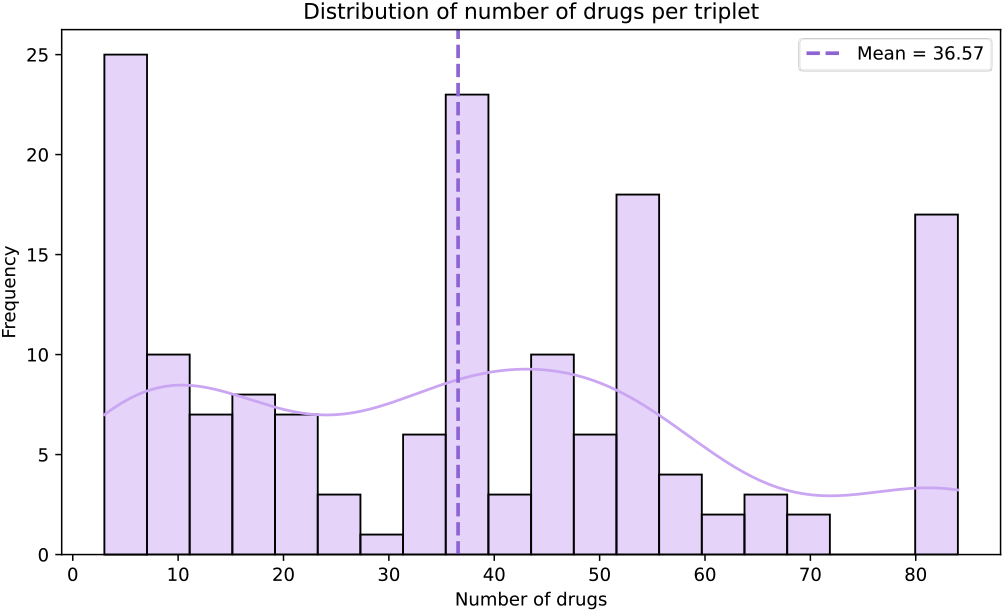
Representation of he number of drugs associated with each triplet combination.

### B. Network Proximity Analysis

As previously described in the Methods section, network *proximity* was calculated to rank candidate drugs based on their topological relationship to the pathway and the disease within the PPI network. For each *Disease* – *Pathway* – *Drug* triplet, we first identified the set of genes shared between the disease of interest (Disease 1) and the associated biological pathway. In parallel, the target genes of each candidate drug were retrieved.

Using these two gene sets, we computed two proximity metrics within the PPI network: the *minimum SPL*, defined as the shortest distance between any gene in the drug target set and any gene in the disease–pathway gene set, and the *closest distance* or *average SPL*, which captures the mean distance between all closest gene pairs across the two sets. Additionally, a Z-score was calculated by comparing the observed *average SPL* to a background distribution generated from 1,000 random gene set permutations of the same size. This Z-score enables the assessment of whether the observed *proximity* between the drug targets and the disease–pathway genes is significantly closer than expected by chance.

The next step in this study involved establishing a ranking of candidate drugs based on the Z-score values obtained from the network proximity analysis. Since lower Z-score values indicate a greater topological proximity between the drug targets and the disease–pathway gene set, candidate drugs were ranked in ascending order of their Z-score for each triplet. After ranking all candidate drugs for each disease– pathway–disease triplet, the top-ranked drug—i.e., the one with the lowest Z-score—was selected as the most promising repurposing candidate for that specific case. **Table II** presents a subset of ten representative triplets, along with their top-ranked candidate drugs. These selected triplets correspond to those with the most negative Z-scores across all analyzed data, highlighting the strongest proximity relationships within the network.

**TABLE 1.** Summary OF THE RESULTS OBTAINED FROM THE NETWORK PROXIMITY ANALYSIS.

|  | Mean | Std | Min | Max |
| --- | --- | --- | --- | --- |
| **Number of shared genes between the disease and the pathway** | 7.5070 | 7.9362 | 1.0000 | 31.0000 |
| **Number of drug target genes** | 4.4351 | 5.1923 | 1.0000 | 51.0000 |
| **Minimum SPL** | 1.9662 | 0.7270 | 1.0000 | 4.0000 |
| **Closest distance (average SPL)** | 2.3885 | 0.5701 | 1.0000 | 4.1666 |
| **Proximity (Z-score)** | -0.0481 | 1.2739 | -4.4487 | 4.8125 |

**TABLE 2.** Top 10 DRUG CANDIDATES RANKED BY NETWORK PROXIMITY.

| Disease | Pathway | Drug proximity average | Top drug | Proximity with top drug |
| --- | --- | --- | --- | --- |
| C0002395 | WP673 | -0.6806 | CHEMBL1336 | -4.4487 |
| C0026769 | WP673 | -0.1856 | CHEMBL553 | -3.9424 |
| C0038356 | WP2363 | -0.8047 | CHEMBL84 | -3.8842 |
| C0278701 | WP2363 | -0.7814 | CHEMBL481 | -3.8832 |
| C0024623 | WP2363 | -0.6935 | CHEMBL1265 | -3.8776 |
| C0278701 | WP2361 | -0.5174 | CHEMBL113 | -3.8274 |
| C1285162 | WP2371 | -0.4526 | CHEMBL1020 | -3.4038 |
| C0005686 | WP2828 | -0.2262 | CHEMBL238071 | -3.2876 |
| C0752347 | WP2371 | -0.3621 | CHEMBL1020 | -3.1653 |
| C0006826 | WP2828 | -0.5196 | CHEMBL806 | -3.1366 |

In the following section, these prioritized drug candidates will be further investigated by integrating transcriptomic signatures, in order to assess their potential functional impact and validate the repurposing hypothesis at the gene expression level.

### C. Connectivity Map – Gene Expression

To further investigate the biological relevance of the top-ranked candidate drugs identified through network proximity analysis, we integrated transcriptomic data from the CMap. The objective was to assess whether these compounds induce significant changes in gene expression when applied to various human cell lines, thereby supporting their potential functional impact on the disease–pathway context. For each of the 48 selected candidate drugs, we analyzed their associated gene expression signatures in CMap. Specifically, we examined whether the transcriptional changes induced by drug treatment showed enrichment in upregulated and downregulated genes, which may reflect modulation of the biological processes associated with the target disease.

The results revealed that all candidate drugs produced detectable alterations in gene expression profiles, with signatures that included both upregulated and downregulated genes across different cell types. On average, each drug induced the upregulation of 2302.67 genes and the downregulation of 1200.22 genes. **Figure 2** provides a heatmap representation of these transcriptomic changes. In this visualization, red indicates upregulation, while blue indicates downregulation of gene expression. The color intensity corresponds to the Z-score values derived from the CMap database, providing a standardized measure of transcriptional perturbation. This integrative analysis supports the biological plausibility of the prioritized candidates and offers an additional layer of evidence for their potential repurposing.

**Fig. 2.**
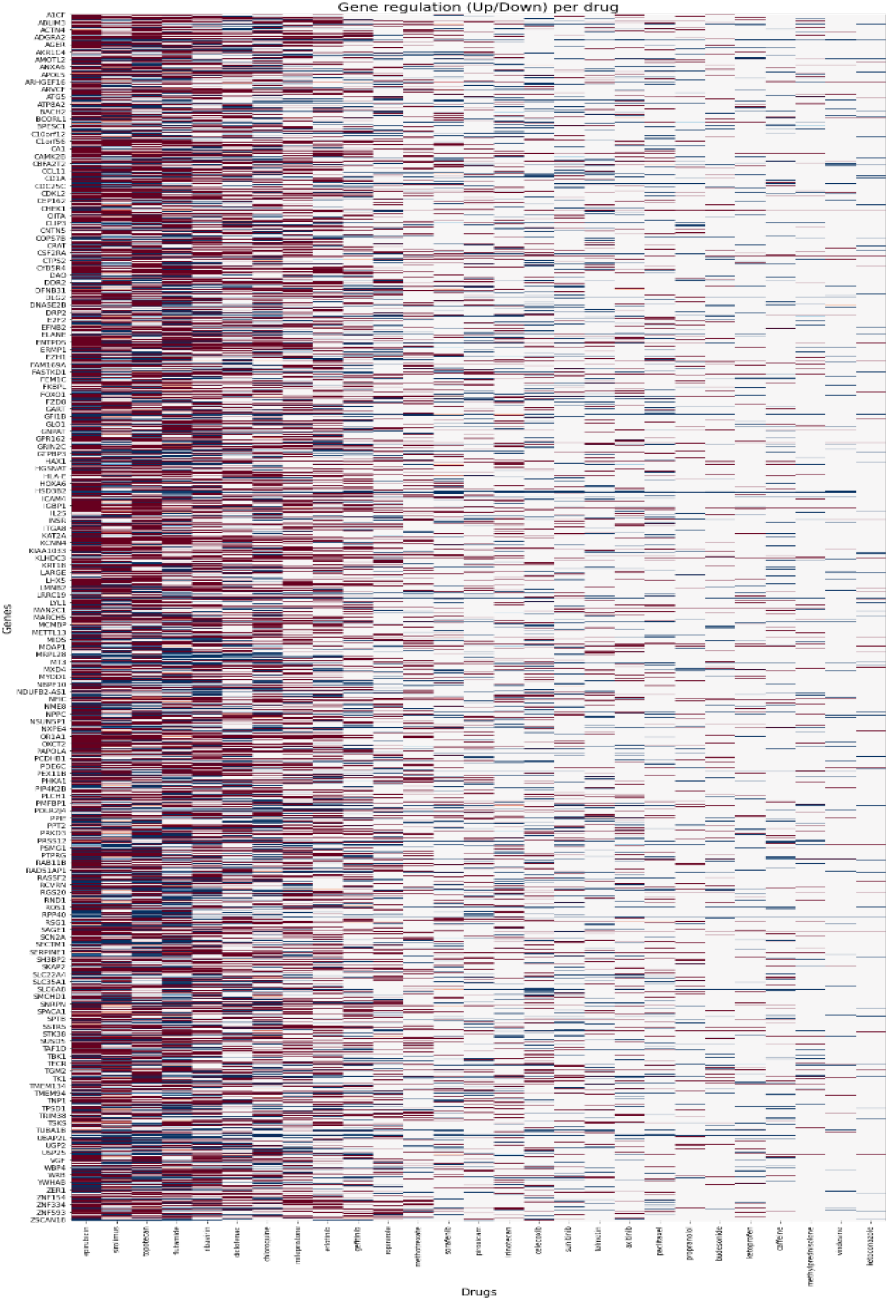
Heatmap of transcriptomic changes, where red indicates upregulation and blue indicates downregulation of gene expression.

Understanding the specific genes modulated by these candidate drugs is crucial, as such modulation can significantly impact therapeutic efficacy and disease progression. For instance, if a drug induces the downregulation of a gene whose overexpression is implicated in the pathogenesis of a disease, this suggests a promising therapeutic interaction. Conversely, upregulating genes that are under-expressed in a disease state can also be beneficial. This concept aligns with previous research indicating that drug-induced gene expression changes can serve as predictive markers for therapeutic outcomes [19]. Furthermore, analyzing drug-induced transcriptional modules has been instrumental in drug repositioning efforts, providing functional insights into how drugs can be repurposed based on their gene expression impact. Such analyses have uncovered cell-type-specific drug responses and highlighted the conservation of drug-induced transcriptional modules across different cell lines and organisms [20].

Given the importance of transcriptional modulation in drug efficacy, our investigation focused on establishing a functional relationship between disease pathways and the transcriptional regulation induced by candidate drugs, as retrieved from CMap. By examining the proximity of candidate drugs to disease-related pathways, we sought to understand how drug-induced gene regulation could impact disease processes.

Figure 3. illustrates this relationship through a network representation showing interactions between drug genes, the drug itself, and the associated diseases. The network highlights the regulation of target genes to explore how they might influence the disease. Only the top-ranked drug candidates for each disease were included, along with the most proximal genes from the drug to the disease pathway. However, not all of these genes were found to be dysregulated in CMap for the respective drug. The figure demonstrates how these regulatory interactions might help inform potential drug repurposing strategies.

**Fig. 3.**
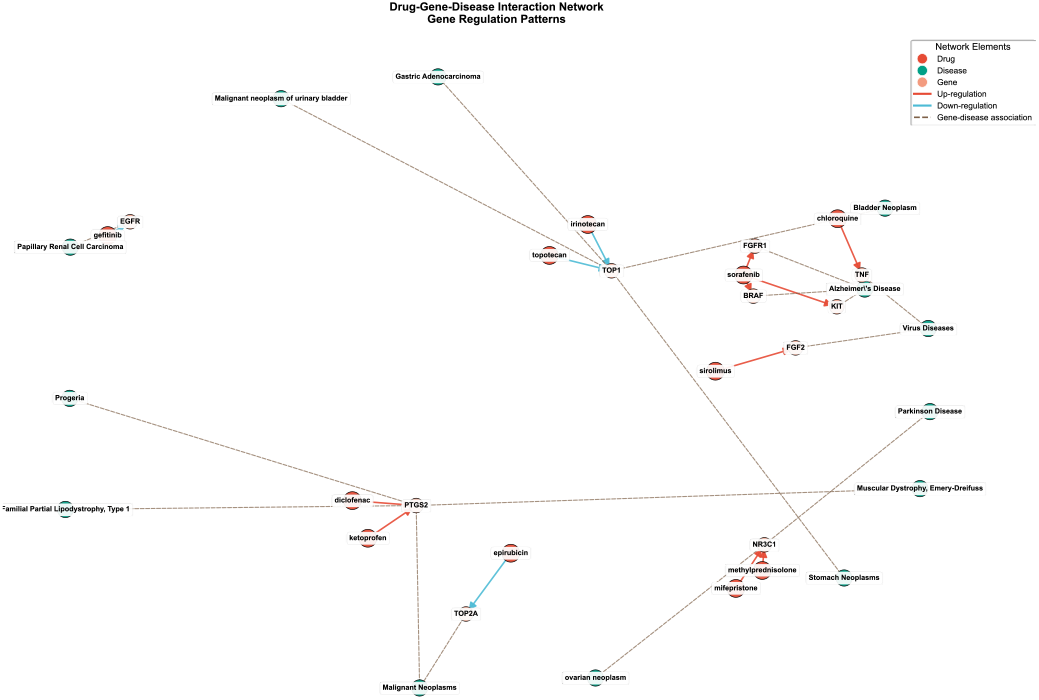
Network showing drug (red)-gene (yellow)-disease (green) relationships. Arrows indicate gene regulation (red=up, blue=down); dashed lines show associations.

### D. Validation

Within this final section, we aim to illustrate a real example derived from our study. To this end, we selected the case with the lowest Z-score across all evaluated triplets, which links Alzheimer’s disease to the drug sorafenib through the ErbB signaling pathway.

Recent studies suggest that sorafenib, a multikinase inhibitor approved for cancer therapy [21], may have previously unrecognized potential in treating Alzheimer’s disease (AD) through modulation of neuroinflammatory responses. In addition to its primary targets (e.g., RAF1, BRAF, VEGFR, PDGFR), some of which are linked to neuronal survival and MAPK/ERK signaling, sorafenib has been shown to affect neuroinflammatory processes that are central to AD pathophysiology. Notably, the study “Sorafenib Modulates the LPS- and Aβ-Induced Neuroinflammatory Response in Cells, Wild-Type Mice, and 5xFAD Mice” demonstrated that sorafenib suppresses the expression of pro-inflammatory markers such as COX-2 and IL-1β in microglial BV2 cells exposed to LPS and reduces astrogliosis in 5xFAD transgenic AD mice challenged with Aβ [22]. These effects were mediated through the inhibition of the AKT/p38 and NF-κB/STAT3 signaling pathways, both implicated in neurodegenerative progression [23]. This evidence supports the hypothesis that sorafenib’s modulation of key inflammatory cascades could attenuate AD-related neuroinflammation. Furthermore, our analysis highlights the interaction between sorafenib’s known gene targets and AD-associated genes via pathway WP673, which involves sirtuin signaling—an axis increasingly recognized for its role in neuroprotection, aging, and metabolic regulation [24]. Collectively, these data suggest that sorafenib’s polypharmacological profile and its influence on disease-relevant pathways make it a promising candidate for repositioning in the context of AD.

## IV. CONCLUSIONS

This study presents a novel pathway-based drug repurposing approach that integrates PPI networks and transcriptomic data to prioritize candidate drugs. By focusing on disease–pathway–disease relationships, we identified meaningful biological links that guided the selection of drugs known to treat one disease (*Disease 2*) as potential therapies for another (*Disease 1*) sharing the same pathway.

Through the computation of network proximity metrics— including shortest path distances and Z-scores—we ranked candidate drugs based on their topological closeness to the disease–pathway module. Further integration with Connectivity Map data enabled us to assess the transcriptional impact of these drugs, revealing that all candidates induced significant gene expression changes, thereby supporting their potential functional relevance. Notably, we found that the genes most proximal to drug targets in the network were frequently transcriptionally regulated by the same drugs, suggesting a strong mechanistic connection between network-based predictions and functional outcomes. Overall, this integrative strategy offers a scalable and biologically informed framework for drug repurposing, enabling the identification of promising candidates with mechanistic support at both the network and transcriptomic levels.

## V. Future steps and limitations

Despite the promising results, several limitations must be acknowledged. First, the analysis relies heavily on the completeness and accuracy of public databases, including PPI networks, drug–gene targets, and transcriptomic profiles from CMap. Incomplete or biased data may affect the reliability of the network proximity and enrichment analyses. Second, the CMap gene expression data are derived from a limited set of human cancer cell lines, which may not fully capture the disease-specific context or tissue heterogeneity relevant to certain drug–disease pairs. Additionally, our current proximity analysis considers static networks and does not account for dynamic regulatory interactions or disease stage-specific changes.

Future steps should focus on expanding the approach to incorporate tissue-specific networks, single-cell transcriptomics, and temporal gene expression dynamics. Incorporating additional omics layers (such as epigenetic data or proteomics) may also improve candidate prioritization. Moreover, in vitro or in vivo validation of top-ranked drug– disease predictions is essential to assess their true therapeutic potential. Finally, incorporating machine learning models could help refine predictions by learning patterns across multiple data types and biological features.

## Acknowledgment

The work is a result of the project “Data-driven drug repositioning applying graph neural networks (3DR-GNN)” that is being developed under grant “PID2021-122659OB-I00” from the Spanish Ministry of Science and Innovation.

## Footnotes

1 https://medal.ctb.upm.es/internal/gitlab/b.otero/pathway-based-dr-network-analysis

